# Climate-Driven Forest Reassembly Follows Divergent Functional Pathways in Cold- and Warm-Adapted Communities

**DOI:** 10.1101/2025.11.18.689113

**Authors:** Ilya Shabanov, Jonathan Tonkin, Andrew Lensen, Julie Deslippe

**Affiliations:** Victoria University of Wellington, School of Biological Sciences, Wellington, New Zealand; University of Canterbury, School of Biological Sciences, Christchurch, New Zealand; Te Pūnaha Matatini Centre of Research Excellence, University of Canterbury, Christchurch, New Zealand; Victoria University of Wellington, School of Engineering and Computer Science, Wellington, New Zealand

## Abstract

**Aim:** Climate change leads to the widespread rearrangement of forests predominantly towards warm-adapted lower-elevation species. However, understanding the context-specific community assembly mechanisms that underlie these changes requires a mechanistic understanding of plant responses. Here, we use functional traits and diversity indices to identify distinct responses of warm- and cold-adapted forest communities.

**Methods:** Our analysis used seven traits, including leaf morphology, size, nutrient content, and foliar isotopes across 3,625 sites over 49 years. We derived functional strategies from community-weighted mean trait values, assessed assembly mechanisms using five multi-trait functional diversity indices, and estimated structural change and turnover rates using abundance-based dissimilarity metrics.

**Results:** Forest composition shifted significantly towards warm-adapted, lowland species with increasing temperature (+0.12 °C/decade) and decreasing precipitation (0.87%/decade). Communities concurrently increased in functional and species richness, but became functionally more similar, indicating niche packing. Disturbance recruitment shifted warm-adapted forests towards acquisitive traits, and increases in anisohydric strategies, reflecting possible effects of CO_2_ fertilisation under drought stress. Cold-adapted forests, in contrast, shifted to conservative strategies and were subject to stronger abiotic filtering. Moreover, changes in nitrogen isotope ratios point towards the importance of ectomycorrhizal associations to persist under climate change. Ecotonal mixed forests showed both responses and experienced the highest turnover.

**Main conclusions:** New Zealand forests follow biome-specific trajectories, characterised by distinct assembly mechanisms and context-dependent ecological responses. While uniform trends obscure these responses, they can be disentangled using a functional diversity framework that incorporates isotopic traits, offering a practical means to detect and interpret community reassembly under climate change.

## 1. Introduction

Forests provide humans with critical resources and services, and influence climate at the global scale (Trumbore et al., 2015). However, forest species composition is changing unpredictably and faster than at any time in the Holocene (Lawlor et al., 2024). Most of the biosphere now experiences novel ecological conditions, where combinations of altered climate, defaunation, and floristic disruption fall outside the historical range and create community structures that are not predictable from past dynamics (Kerr et al., 2025; Radeloff et al., 2015). Reduced tree longevity (Locosselli et al., 2020), altered reproductive phenology (Flores et al., 2023), heightened vulnerability to pathogens (Wyka et al., 2018), and intensified pressure from insect outbreaks (Pureswaran et al., 2018) collectively diminish productivity (Mao et al., 2022), carbon sequestration capacity (Carnicer et al., 2019), and resilience (Oliver et al., 2015). These accelerating trends and ecological novelty create an urgency to understand the mechanisms of biodiversity rearrangement, to maintain the health of Earth’s forests, and to preserve its biodiversity.

Observations of biodiversity rearrangement alone, so far, have not provided sufficient grounds to predict these accelerating changes (Rubenstein et al., 2023). For instance, in some places, climate warming has induced expansions of warm-adapted species (Gottfried et al., 2012) (thermophilization) and a concomitant decline of cold-adapted species (Rumpf et al., 2019). By contrast, biodiversity can remain unchanged (Dornelas et al., 2014; Vellend et al., 2013), increase (Montràs-Janer et al., 2024; Steinbauer et al., 2018), or decrease (Pereira et al., 2024), even in strongly affected Arctic ecosystems (Cotto et al., 2017; García Criado et al., 2025). The speed of biodiversity change depends on the climate (Fadrique et al., 2018; Stephenson & Mantgem, 2005), but is rarely quantified. Underlying these conflicting patterns in the rearrangement of forests are context-dependent mechanisms (Lawlor et al., 2024; Lenoir & Svenning, 2015) interacting with hitherto unobserved, novel climatic conditions (Staples et al., 2022). Moreover, forests exhibit response lags due to long generation times (Breshears et al., 2008), which can be overlooked in comparatively small or short-term datasets (Komatsu et al., 2019;Valdez et al., 2023). Reliable predictions of biodiversity rearrangement, therefore, require a mechanistic understanding of plant-environment interactions, inferred from larger and longer datasets.

Leveraging plant functional traits in models to predict plant responses to environmental change offers a more mechanistic approach to plant-environment relationships (Funk et al., 2017). However, while relationships between single traits and environmental gradients are well established (Wieczynski et al., 2019), they lack explanatory power and fail to capture the complexity of interacting drivers (Green et al., 2022). Trait syndromes, on the other hand, reflect ecological strategies that allow species to succeed in an environment (Laughlin, 2023) and enable predictions of which species will thrive there in the future (Henn et al., 2024). For instance, specific leaf area (SLA) and leaf nitrogen content concurrently increase with temperature and decrease with elevation, suggesting an adaptation to harsher, nutrient-poorer environments (Homeier et al., 2021). Conversely, lower SLA with higher stem density is associated with higher drought tolerance (Greenwood et al., 2017). Capturing the dynamics of trait syndromes at the community scale holds the key to understanding forest responses and to better forecast changing biodiversity patterns under previously unobserved conditions (Kerr et al., 2025).

The ecological insights derived from trait data are constrained by the traits selected. Measured traits should therefore align with the trait syndromes and functional strategies most relevant to the climatic context (Hagan et al., 2023). For instance, much of research has focused on easily measured leaf, seed, and stature traits (Green et al., 2022), which capture broad plant strategies along the r-K spectrum (Díaz et al., 2022), but are less suitable for explaining water use strategies critical in arid environments (L. Li et al., 2015). In contrast, underutilised traits such as nitrogen-15 (Δ^15^N) and carbon-13 (Δ^13^C) isotopes provide more mechanistic insight into nutrient acquisition strategies, water-use efficiency (Dawson et al., 2002),(Dawson et al., 2002), and plant-microbe associations (Song & Zhou, 2021), while also informing productivity (Luo et al., 2009) and nutrient limitation (McLauchlan et al., 2010) in ecosystems. Accurately forecasting plant community responses to environmental change, therefore, requires an understanding of a broad suite of traits encompassing diverse functional strategies.

The relevance of traits and functional strategies depends on the environmental constraints faced by the community (Lebrija-Trejos et al., 2010). For instance, strategies favouring rapid resource acquisition are disadvantageous in harsh environments that favour slow-growing, stress-tolerant species (e.g. alpine tundra; (Spasojevic & Suding, 2012) but advantageous in resource-rich environments where competition shapes community assembly (Grime, 1977). These assembly processes can be inferred from relationships between functional richness (FRic), the range of strategies present, and functional dispersion (FDis), the degree of trait differentiation (Laliberté & Legendre, 2010; Lamanna et al., 2014). Decreasing FRic and increasing FDis suggest selection for extreme but distinct strategies, revealing strong abiotic filtering. Conversely, increasing FRic and stable FDis suggest differentiation of species into distinct niches. Strong abiotic filtering excludes species with traits unsuited to local stressors, producing communities of functionally similar, stress-tolerant species (Le Bagousse-Pinguet et al., 2017). In contrast, niche complementarity among diverse traits promotes efficient resource use, multifunctionality, and greater resilience (Gross et al., 2017; Searle & Chen, 2020). Functional diversity patterns should therefore be considered alongside trait analyses as proxies for assembly mechanisms, enabling insight into both the traits favoured and the long-term structure of communities.

Here, using 49 years of vegetation data from 3,625 forest plots across New Zealand and nine functional traits within a functional diversity framework, we aim to understand the mechanisms behind biodiversity rearrangement in forest communities. To separate broad climatic effects from biome-specific dynamics, we first partition communities into cold-adapted, warm-adapted, and mixed ecotonal forest types and identify the community assembly mechanisms constraining the ecological strategies in each biome. Next, we identify which functional strategies drive shifts in these communities to reveal community-specific adaptations to novel climatic conditions. Lastly, we aim to quantify the tangible ecological consequences of these processes by estimating the rate of turnover within these communities. Together, these aims provide a basis for understanding multiple pathways of forest reassembly and for anticipating long-term biodiversity trajectories under accelerating climate change.

## 2 Methods

### 2.1 Species Occurrence, Abundance Data, and Forest Types

We utilised forest plots in New Zealand’s national forest inventory data (NVS, (Wiser et al., 2001)), which were measured using two different protocols: RECCE and DBH. The RECCE protocol (Hurst, 2022) contained categorical cover estimates for every height tier category (1 through 6, for individuals > 25m down to < 60cm), collapsible into a single cover estimate for each species and plot (Fischer, 2015). The sum of these estimates was normalised to 1, yielding relative canopy cover values for each species The DBH protocol contained individual adult tree counts and their diameter at breast height (DBH). The sum of all individual DBH values per species in a plot was used as an abundance metric. As ∼25% of plots were measured using both protocols, we were able to compare the two abundance metrics. They showed a strong correlation of R^2^=0.68, enabling us to combine plots from both protocols into a single dataset containing a list of species and their relative abundance in the plot.

Finally, we assigned one of three forest types (Beech, Podocarp (which also included broadleaved species), or Mixed) to each plot using the 25m Basic Ecosystem layer from New Zealand’s Landcare Research database (Landcare Research, 2024), as it is the only nationwide forest vegetation classification dataset for existing rather than historic vegetation. The forest type where a species was most abundant was declared its primary forest type. However, most species were present in all three forest types with varying abundance. Shrublands were excluded, as they contained young successional stands of regenerating forests, which are the result of land-use change, rather than climate change and follow different community assembly mechanisms than forests (Subedi et al., 2019).

### 2.2 Trait Data

We collected and compiled trait data on seven traits: Specific Leaf Area including the petiole (SLA), stem specific density (SSD), leaf nitrogen content per unit mass (Nmass, mg/g), Δ^13^C carbon isotope (in ‰), Δ^15^N nitrogen isotope (in ‰), diaspore mass (in mg) and plant height (in m). We collected trait data on the 78 most abundant and locally available tree, shrub, and vine species in the Wellington (New Zealand) area from September 2024 to February 2025. Collection, handling, and measurement followed the protocol in (Pérez-Harguindeguy et al., 2013). Additionally, we downloaded all available traits of the New Zealand flora from the TRY database (Kattge et al., 2020) and the Ecotraits database (*EcoTraits Landcare Research*, 2005). Measurements from all three sources were combined by taking the average trait value of all entries. To reduce errors, we discarded species with more than three out of seven trait values missing. This resulted in a final set of 139 species we could use for our analysis.

Missing trait values (ca. 22%, ***Fig. S2*** B) were imputed using Multiple Imputation by Chained Equations (MICE) with taxonomic eigen vectors (Debastiani et al., 2021) and the miceForest Python implementation (Wilson et al., 2023). This algorithm exhibits superior performance compared to other contemporary methods (Penone et al., 2014). As MICE is not deterministic, we re-ran the imputation algorithm five times and averaged the imputation results. The trait distributions before and after imputation were minimally different and didn’t significantly change the correlations between the traits (***Fig. S2*** A and C), validating the imputation approach. The average imputation error was 10.68% of the standard deviation of the trait. Details of the process and error metric are described in Section 2 of the supplementary materials.

### 2.3 Environmental Variables

We selected a set of = *16*variables from NZEnvDS (*New Zealand Environmental Data Stack (NZEnvDS) - Manaaki Whenua - Landcare Research DataStore*, n.d.), which included topographic (e.g., elevation), edaphic (e.g., pH), and climatic variables (e.g., mean annual temperature), and determined their values at the plot locations. This dataset was preferred due to its high spatial resolution (100m).

To assess the climate trend, we obtained monthly rainfall and minimum and maximum temperatures (albeit at a lower 1km resolution) from the HOTRUNZ dataset (Etherington et al., 2022) and converted them into BioClim variables (O’Donnel & Ignizio, 2012). For each plot, we approximated the time series of each variable over the years (1970-2019) with a linear function and used its slope as a measure of climate change at this location. Positive values indicate an increasing trend of this climatic variable over time. To assess whether climate change affects the forest types differently, we compared the slope distributions between the plots belonging to the three forest types and computed Cohen’s D as a measure of difference in climate change impact between them (Cohen, 2013). Cohen’s D measures the difference in means in relation to the difference in standard deviations. Values are commonly binned in small effects: 0.2-0.5, medium effects: 0.5-0.8, and large effects: above 0.8.

### 2.4 Community Weighted Mean (CWM) trait values

The community-weighted mean trait value is a plot-level variable representing the prevailing trait value of the community at the plot scale used to predict ecosystem functions, such as primary productivity (Violle et al., 2014). It is calculated using the average of species trait values () weighted by the abundance of the species () in a plot :,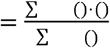. We calculated CWMs only for plots with five or more species. We tested for CWM changes over time to assess functional strategies of communities (Aim 2).

### 2.5 Community Environmental Index

The average thermal preference of all species in a community, weighted by their relative abundance, is referred to as the community temperature index (CTI, (Devictor et al., 2008)).

Extending this definition to other plot-level variables (climatic, topographic, and geographic), we defined the community environmental indices (CEI). The CEI represents the average of the species’ environmental preference (SEP) weighed by their abundance in the plot : () and is defined as 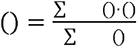 . Where 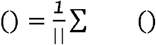 is the mean plot-level environmental variable (e.g., temperature or elevation) across all plots where the species has been observed The () is a plot-level variable that can be interpreted as the environmental preference of the entire community inside a forest plot defined by its species. The CEI for temperature is thus identical to the CTI. We evaluated CEI trends over time using linear mixed models (see Section 2.9) to probe for effects of thermophilization (increasing CEI for mean annual temperature), invasion by lowland species (decreasing CEI for elevation), and others (***Fig. S3***).

### 2.6 Functional Diversity Indices

To assess community assembly mechanisms (Aim 1), we computed five functional diversity indices for each plot following (Pla et al., 2012): Functional Richness (FRic), functional divergence (FDiv) and functional evenness (FEve) (Villéger et al., 2008), the mean nearest neighbour distance (MNND) in trait space (Swenson & Weiser, 2014), and functional dispersion (FDis) (Laliberté & Legendre, 2010). FRic is calculated as the volume of the convex hull of species’ T trait values for each plot and therefore requires at least T + 1 species to be present (here, T=7). MNND is calculated as the average Euclidean distance in trait space between each species and its nearest neighbour. These indices are used to quantify the size and packing of trait space. A large FRic value indicates a community with many diverse trait values and therefore correlates well with species richness (Swenson & Weiser, 2014). MNND indicates how similar or packed these species are in trait space, with low values indicating the clustering of functionally similar species. FEve quantifies how evenly species abundances are distributed along trait axes, while FDiv reflects how far species deviate from the centre of trait space, emphasising the presence of functionally extreme species. FDis captures the mean distance of individual species to the community centroid, weighted by relative abundance, thus representing trait dispersion within the community. We transformed to = (*1*+) to normalise its distribution and facilitate linear regression.

### 2.7 Rate of Turnover: Index of Dissimilarity change per Year(IoDpY) for species

To assess aim three, we modified the Index of Dissimilarity (Taeuber & Taeuber, 1976), IoD) which measures what proportion of individuals (or basal area in our case) would need to move to achieve the observed species rearrangement. A value of 1 indicates that all individuals would need to relocate, while a value of 0 means no change. We used IoD as a metric of turnover for single species. It was determined by using the abundances of species in remeasured plot pairs (^0^() and ^1^()) and their sums across all plots (e.g. ^1^() = ∑ ^1^()). Giving the index: 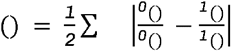. Since IoD would be expected to be higher if the remeasurement interval is increased, we calculated an IoD per year using the following scheme:

1. Binning all plot pairs based on their remeasurement interval ()
2. Computing the IoD for each subset (,)separately
3. Combining the IoDs divided by and weighed by the number of plots for each interval | |:

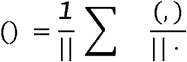

The resulting metric is independent of the remeasurement interval. IoDpY can be interpreted as the percentage of a species currently occupied basal area that needs to shift per year to match the observed species turnover. However, it does not inform whether a species expands or contracts its range.

### 2.8 Rate of Directional Change: Estimating plot occupancy for species over time

To estimate whether a species has increased its plot occupancy over time, we calculated the ratio of plots containing this species to the total number of plots measured in that year, yielding a single plot occupancy value of 0-1 for each year (omitting years where the species was not detected). This is done for each forest type separately. We then ran a linear regression to determine whether plot occupancy had a significant positive trend (species becomes more common) or negative trend (species becomes less common). We used the slope of the linear regression to indicate the rate of change in occupancy, interpreting only slopes statistically significantly different from zero. This allows us to qualitatively compare occupancy trends of species across forest types. As the location of the plots is not considered, the rate of occupancy change does not differentiate a species becoming more common within its geographical range from an expansion of the species’ range. In both cases, a positive rate of occupancy change indicates an increase in individuals of this species in response to climate change.

### 2.9 Linear regressions and statistical testing

For temporal trends analysis (CEI, diversity indices, species richness, and CWMs of traits), we fitted linear mixed-effects models using the statsmodels Python package (Seabold & Perktold, 2010), with plot ID included as a random intercept to account for repeated measurements of the same plot. Fixed effects (e.g., Year) were tested using Wald tests, assuming an asymptotic normal distribution of the test statistic. Parameters were estimated via restricted maximum likelihood. For estimating the rate of change, a regression was run on estimates per year, using ordinary least squares regression with the same significance testing.

## 3 Results

### 3.1 Widespread shifts towards warm-adapted, low-elevation species

Community environmental indices for elevation, latitude, mean annual temperature, slope, and precipitation (**Fig. 1**, complete list in ***Fig. S3***) for plots of individual forest types showed a coordinated response. CEIs of all forest types shifted towards lower elevations, higher mean annual temperatures, and lower latitudes (away from the poles), indicating a thermophilization of communities from lower elevations, by more warm-adapted higher-latitude species (**Fig. 1**B). Notably, only warm-adapted podocarp forests exhibited community shifts towards more drought-adapted low precipitation species.

**Figure 1.**
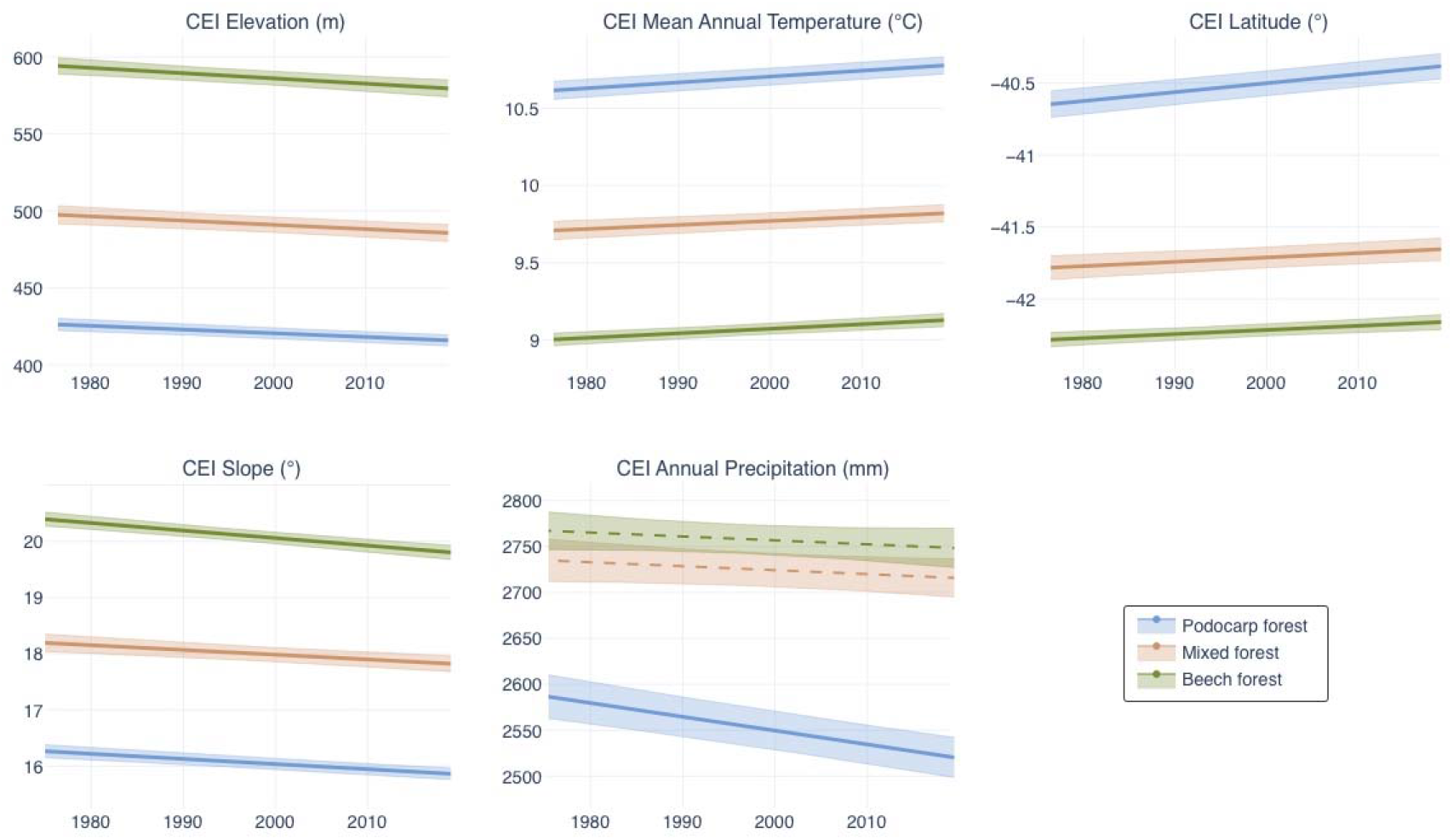
Temporal trends of community environmental indices (CEI, reflecting the environmental conditions favoured by the species present in a plot) for five plot-level variables over time for the three studied forest types (indicated by colour). Significant relationships are displayed as solid lines (insignificant ones are dashed); 95% confidence intervals are shaded.

### 3.2 Niche packing and differential traits in functional divergence

In all three forest types, taxonomic and functional richness (FRic) and dispersion (FDis) increased simultaneously, accompanied by a decrease in MNND, indicating broader niches with more tightly packed species. Beech forests generally showed less change over time. FDiv increased in podocarp forests, but declined in beech forests, with FEve showing no significant changes (**Fig. 2**).

**Figure 2.**
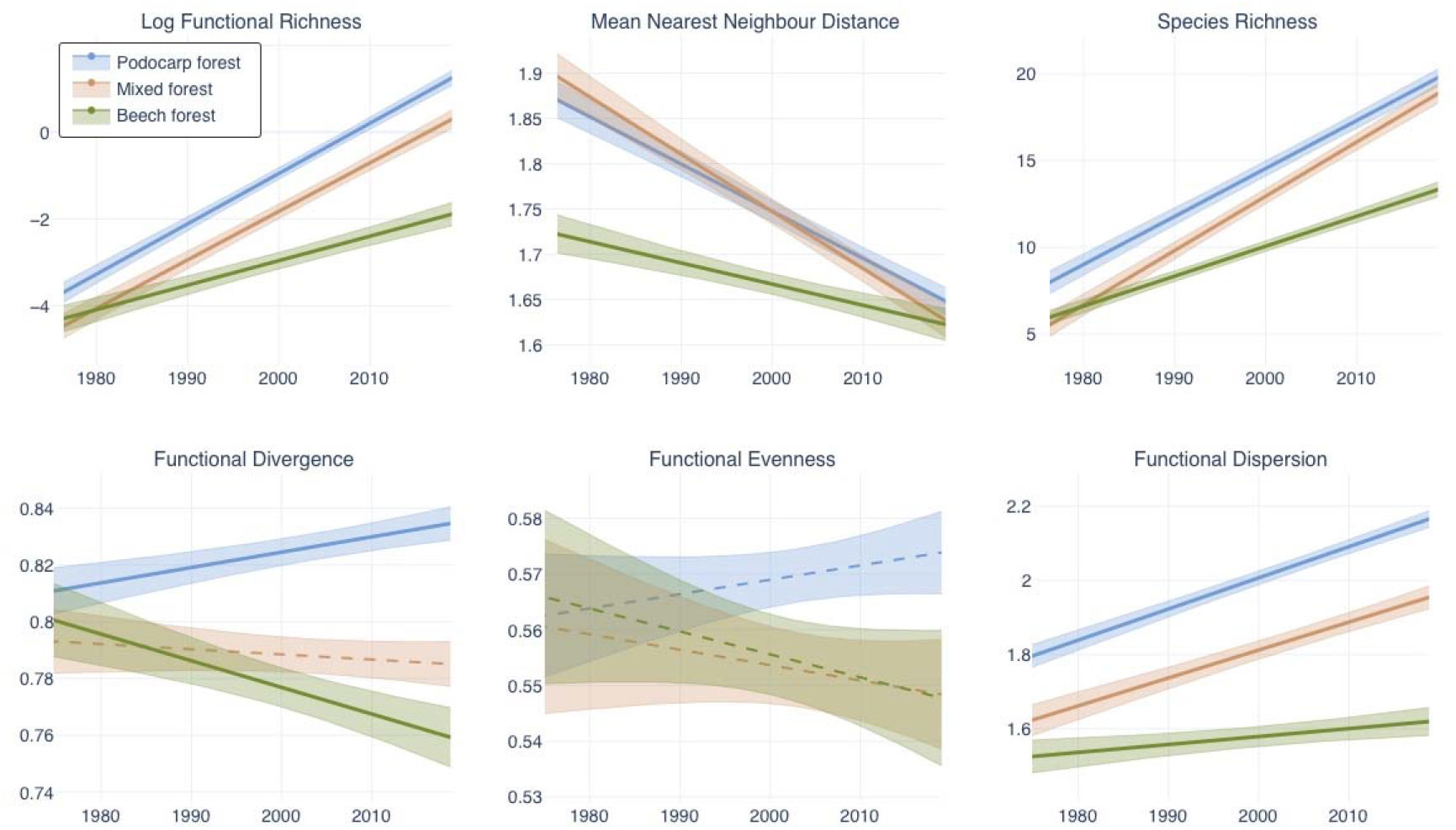
Temporal trends of multi-trait functional diversity indices and species richness for the three studied forest types (indicated by colour). Significant relationships are displayed as solid lines (insignificant ones are dashed), 95% confidence intervals are shaded.

### 3.3 The rate and identity of species change and turnover differ by forest type

Across forest types, a similar number of species showed non-zero rates of change (28-34, **Fig. 3**A, left). However, weighing these species by the cover they occupy revealed that in podocarp and mixed forests, the shifting species are more widespread, leading to larger changes overall (**Fig. 3**A, right). The rate of turnover was highest in mixed forests (**Fig. 3**B). The rate of turnover was only weakly correlated across forest types (R^2^ = 0.06-0.19), indicating that unique species changed in each forest type. We also assessed the primary forest type of species with non-zero rates of change (**Fig. 3**C). In podocarp forests, podocarp forest and mixed forest species expanded, but beech species predominantly contracted. This was contrasted by mixed and beech forests, where all species predominantly expanded their ranges, indicating a differential dynamic in podocarp forests. Importantly, mixed forests also showed a higher rate of change overall.

**Figure 3.**
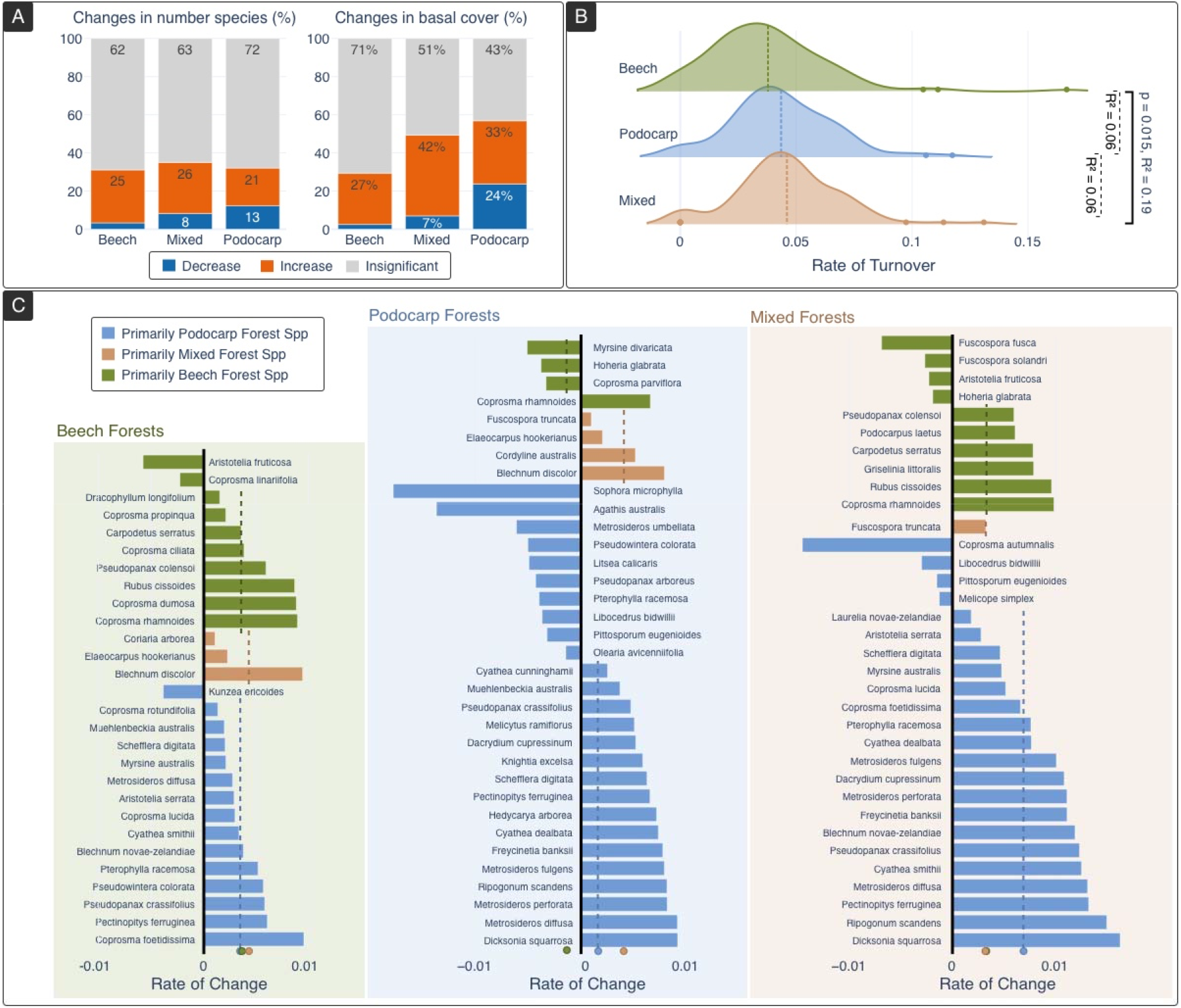
Significant changes for individual species in the rate of directional change (i.e., slope of the occurrence probability over time) and turnover (i.e., percent of basal cover needing to shift per year to match observed turnover) by forest type. (A) Left: Number of species experiencing significant directional changes. Right: Total cover occupied by these species. (B) The rate of turnover for all species found in plots of a forest type. As species often occur in all forest types, their turnover rates can be correlated. Correlation value provided in brackets. (C) Rate of directional change for individual species within each forest type (columns), only significantly changing species are shown. Colours indicate a species’ primary forest type (i.e., the forest type where it is most abundant). Means of bars of the same colour are indicated by a dashed line and a circle on the x-axis.

### 3.4 Functional shifts in forests differ by forest type

The forest types showed marked differences in trait CWMs over time (**Fig. 4**). Podocarp forests decreased in CWMs for Δ^13^C and SSD, while beech forests decreased in leaf nitrogen (NMass), specific leaf area (SLA), and Δ^15^N, indicating distinct functional adaptations. Mixed forests showed similarities with both other forest types, leaf nitrogen and Δ^15^N decreased similarly to beech forests, and SSD similarly to podocarp forests. Additionally, seed mass exhibited a significant decline over time.

**Figure 4.**
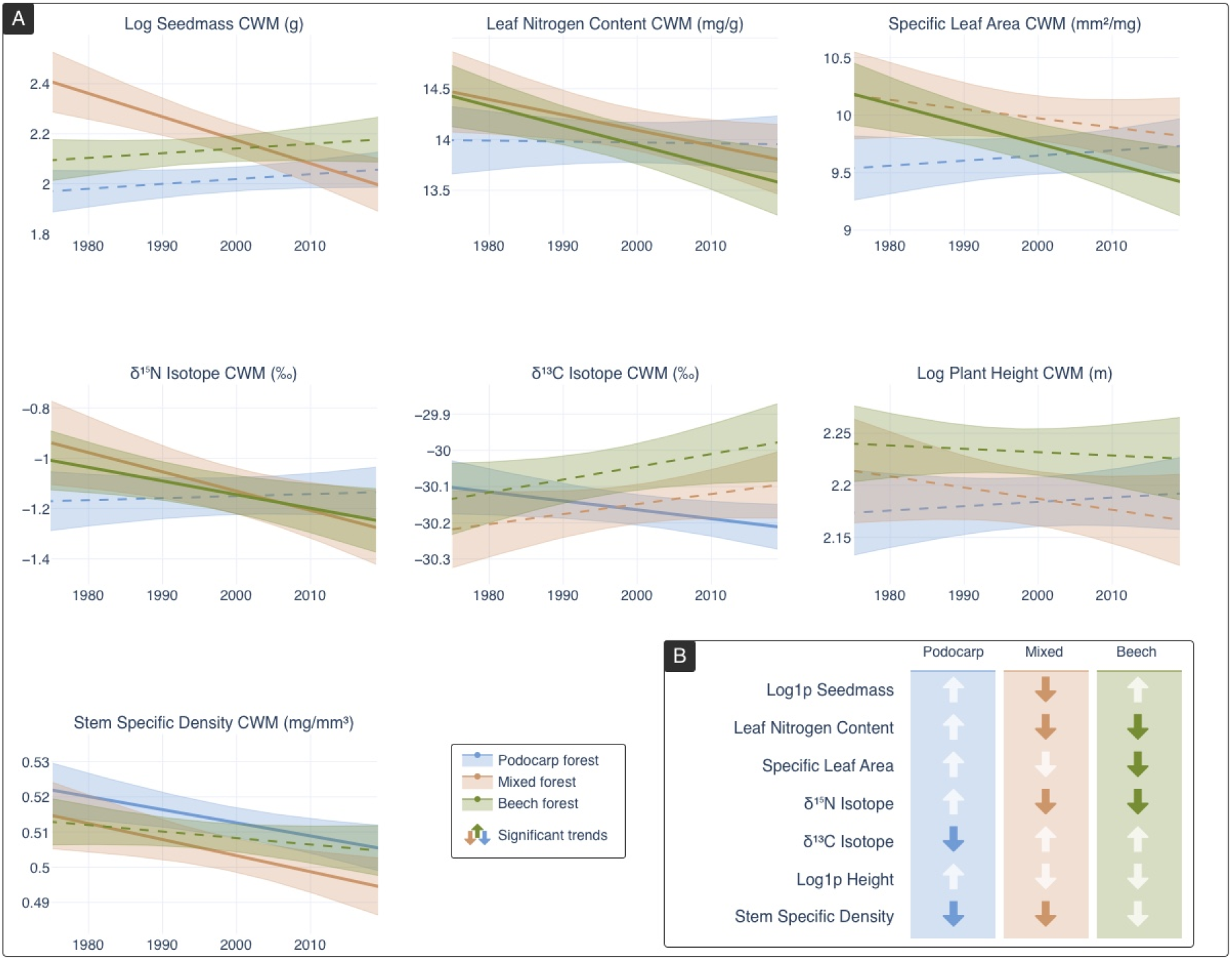
(A) Temporal trends of functional trait CWM for the three forest types (indicated by colour) over time. Significant relationships are displayed as solid lines (insignificant ones are dashed), 95% confidence intervals shaded. (B) Summary of trends observed in panel (A), insignificant shifts are indicated by white arrows.

## 4 Discussion

Our study revealed that climate-driven reassembly of forests along an expansive latitudinal gradient has been structured by contrasting community assembly mechanisms (Aim 1), expressed through divergent functional strategies (Aim 2), and characterised by distinct rates of species turnover (Aim 3). New Zealand’s climate has become warmer and drier (***Fig. S4***), leading to thermophilisation by low-elevation, warm-adapted species (**Fig. 1**), higher local species richness, and pronounced functional changes (**Fig. 4**). Cold-adapted beech forests were shaped primarily by abiotic filtering, producing more conservative trait syndromes characterised by hardier, less nutrient-dense leaves, whereas warm-adapted podocarp forests exhibited more acquisitive strategies with lower Δ^13^C and stem density, reflecting responses to drought and CO_2_ enrichment. Ecotonal mixed forests showed the highest turnover rates and a combination of both functional shifts. Although it is known that species turnover is not consistent across elevation (Fadrique et al., 2018) and latitude (Freeman et al., 2021), our study revealed that distinct community assembly mechanisms cause these differences and result in biome-specific ecological strategies that can be readily revealed using a broader set of traits within a framework of functional diversity.

Over the past half-century, New Zealand has experienced a warming of 0.12°C per decade (***Fig. S4***A), consistent with global estimates of 0.13 °C ± 0.03 °C (Pacifici et al., 2017). This resulted in increased species richness (**Fig. 2**), which has also been observed globally (Steinbauer et al., 2018), but contradicts widespread expectations and forecasts of biodiversity decline (Isbell et al., 2023; Pereira et al., 2010; Rumpf et al., 2019). This phenomenon is likely because only a subset of species can track the changing climate, while others persist or react slowly (Dullinger et al., 2012; Svenning & Sandel, 2013), leading to an initial accumulation of species, but delayed extinctions (Tilman et al., 1994). Over time, a substantial loss of biodiversity can occur (Engler et al., 2011). Assessing species persistence from functional traits, however, remains a challenge as it depends, among others, on life span (de Witte et al., 2012), presence of persistent seed banks (Ouborg & Eriksson, 2004), phenotypic plasticity, standing genetic variation, and evolutionary potential (Cotto et al., 2017; Valladares et al., 2014). Further data and monitoring will help contextualise these patterns within long-term biodiversity trajectories.

Over time, species and functional richness (FRic) increased while trait differences (MNND) decreased (**Fig. 4**), indicating that the functional space is becoming more densely packed with functionally similar species, a pattern consistent with niche packing driven by biotic interactions (Swenson & Weiser, 2014). Concurrently, rising functional dispersion (FDis) reflects more widely scattered traits, reflecting alternative strategies under environmental stress during climate-driven reassembly (Griffin-Nolan et al., 2019; Spasojevic & Suding, 2012). These patterns suggest community assembly balances fine-scale biotic sorting and broader-scale abiotic filtering. Notably, functional divergence (FDiv), a measure of the prevalence of peripheral trait values, increased in podocarp forests, while it decreased in beech forests. This pattern indicates that novel trait values of migrants are more prevalent in podocarp forests, likely because cold-adapted beech forests respond more slowly (Dullinger et al., 2012; Svenning & Sandel, 2013). Importantly, a prevalence of peripheral trait values reflects stress-adapted functional strategies associated with lower primary productivity (Y. Li et al., 2022). While successional dynamics influence these patterns, the stable functional evenness observed here is atypical for successional trajectories (van der Sande et al., 2024). Thus, while species richness increased across all forest types, the pathways of community assembly may differ according to forest type, possibly reflecting contrasting functional strategies and assembly processes between cold- and warm-adapted communities. Podocarp forest communities shifted toward species associated with lower precipitation niches (**Fig. 1**), presumably due to increased precipitation seasonality (***Fig. S4***). However, the observed trait trends of lower Δ^13^C and SSD (**Fig. 4**) indicate a functional transition toward acquisitive species with low water-use efficiency (Farquhar et al., 1989) and may reflect CO_2_ fertilisation, where enhanced carbon uptake alters the C:N stoichiometry, potentially intensifying nutrient limitation (Domec et al., 2017). This imbalance may favour anisohydric species that maintain stomatal conductance under drier conditions, enhancing photosynthetic rates despite water stress (Wang et al., 2021). Although data on anisohydry levels in the New Zealand flora are limited, ferns are generally more anisohydric than overstory species (Hollinger, 1987), and expand rapidly in our dataset (see *Blechnum, Cyathea*, and *Dicksonia*spp in **Fig. 3**C), lending support to this hypothesis. However, the performance advantage of anisohydric species is typically confined to moderate drought and may become a liability under prolonged water deficits (Sade et al., 2012). Given that species with lower SSD are generally more drought-sensitive (Bruelheide et al., 2018; Greenwood et al., 2017), the negative SSD trend may instead originate from mesic forest patches, where increased disturbances promote the recruitment of acquisitive, early-successional, or gap-specialist species with lower SSD (Henn et al., 2024). This observation is consistent with projected functional shifts under climate change (Chain-Guadarrama et al., 2018). Together, these findings suggest that compositional change reflects multiple ecological strategies, rather than a singular response to abiotic filtering.

Cold-adapted forests, dominated by beech species, exhibited significantly slower turnover rates (**Fig. 3** B). This ecological inertia is well-documented (Cotto et al., 2017; De Frenne et al., 2013; Dullinger et al., 2012) and experimentally confirmed (Cox et al., 2024). Cold-adapted beech forest species declined within warm-adapted podocarp forests (**Fig. 3**C), mirroring French temperate forests (Borderieux et al., 2024) and suggesting local extinctions of cold-adapted species, rather than expansion of warm-adapted species, as drivers of thermophilization. Functionally, beech forests shifted towards more conservative strategies characterised by declining SLA and Nmass (**Fig. 4**), suggesting abiotic filtering. This narrowing of functional strategies is further supported by a modest increase in FRic and near-constant FDis (**Fig. 2**). Interestingly, foliar Δ^15^N CWM declined, indicating possible expansions of species with ecto- and ericoid mycorrhizal fungal associations. These associations reduce foliar Δ^15^N (Hobbie & Hobbie, 2008), are common in cold-adapted forests (Soudzilovskaia et al., 2017), promote biomass under elevated CO_2_ (Bennett & Classen, 2020), and buffer trees against abiotic stress (Usman et al., 2021). Their increase suggests an eCO_2_-driven shift in C:N stoichiometry that heightens nitrogen limitation and favours species making greater use of these symbioses (Cambron et al., 2025). Ultimately, the inertia of cold-adapted beech forests, coupled with emerging nutrient constraints, renders them particularly vulnerable to climate change.

The mixed forest ecotone exhibited the highest turnover rate (**Fig. 3**B) and functional diversity changes (**Fig. 2**), likely due to its position between cold- and warm-adapted communities. The observed functional shifts in mixed forests (**Fig. 4**B) reflect a blend of the trait syndromes found in both beech and podocarp forests, suggesting a mixture of their assembly mechanisms. Exposure to contrasting climate responses leads to frequent shifts in species dominance (Peters, 2002) and ecological novelty, characterised by high rates of reassembly (Abbasi et al., 2024). The decrease in seed mass unique to mixed forests likely reflects the expansion of lightweight-diaspore ferns (e.g., *Cyathea, Blechnum, Dicksonia spp*in **Fig. 3**C), which thrive under disturbance (Arens & Baracaldo, 1998), strongly influence community assembly in New Zealand (Brock et al., 2018; Gaxiola et al., 2008), and may therefore act as early indicators of ecological novelty. Ecotones may facilitate high turnover by integrating distinct assembly processes from adjacent cold- and warm-adapted communities, thereby serving as hotspots of ecological novelty.

Although this study benefits from a uniquely large and long-term forest dataset significantly surpassing suggested benchmarks of thousands of sites over decadal intervals (Valdez et al., 2023), several limitations warrant consideration. First, our analysis was restricted to woody species, which do not represent total plant biodiversity and whose responses may not be generalisable beyond forests. Second, we were unable to account for disturbance regimes or land-use change, as spatially explicit information on disturbance currently does not exist for New Zealand forests. Third, trait data imputation, although methodologically robust with reasonably low error rates, may still introduce uncertainty, particularly for rare traits such as Δ^15^N and Δ^13^C. Lastly, our approach ignores species’ intraspecific trait variation in response to the environment. Since our study spans a broad spatial range (subtropical to montane forests, over ca. 1600km), we expect interspecific to vastly exceed intraspecific trait variation(Albert et al., 2011; Lajoie & Vellend, 2015). Despite these limitations, the breadth and consistency of our findings suggest they do not compromise the main conclusions.

## Conclusion

Climate change is reshaping forest communities through divergent ecological pathways, even where trends in species richness and thermophilization appear superficially uniform. Cold-adapted forests are subject to strong abiotic filtering and shift toward conservative, stress-tolerant strategies, whereas warm-adapted forests show higher competition and reassemble more rapidly through acquisitive strategies. This distinction and relation to physiological mechanisms becomes apparent only by leveraging multi-trait functional diversity indices and less-studied traits, such as Δ^13^C and Δ^15^N. Moreover, isotope-based traits suggested an increase in species with mycorrhizal associations in cold-adapted forests, due to a stoichiometric C:N imbalance caused by elevated CO_2_ and limiting available nutrients. In warm-adapted forests, they revealed a shift towards anisohydric species that maintain stomatal conductance in dry conditions as a possible reaction to increasing CO_2_ under decreasing precipitation. Our findings underscore the need for trait-rich datasets to disentangle context-dependent responses and anticipate long-term ecosystem trajectories (Thuiller et al., 2013). Concomitantly, we demonstrate how imputation methods can aid this development by substantially and reliably increasing the number of studied species (Penone et al., 2014). As climate novelty accelerates globally, functional trait approaches that integrate physiological and ecological dimensions provide a scalable framework for understanding and forecasting the future of forest biodiversity.

## Data availability statement

Trait Data and imputation code is available for peer review under the anonymous link: https://zenodo.org/records/16938264?token=eyJhbGciOiJIUzUxMiJ9.eyJpZCI6IjAxYzVkZmM4LWJiZmUtNDk4ZS04ZmIxLTlkY2ZlNmQ0MzQ4NSIsImRhdGEiOnt9LCJyYW5kb20iOiJjNmY0YWZiOTU5MTIyMTBkOTk2MDNhNzg5YjhkM2RiZCJ9.OaeZxaexPgVWYoN3tBKVogyDWRoP1rLT9O-25DFL3fh8UtWudSuXVtvu5Rb1jqbWIzBxYQYQ6zmFh3TmCanZtA

