## Supplementary Materials for "Climate-Driven Forest Reassembly Follows Divergent Functional Pathways in Cold- and Warm-Adapted Communities"

### 1 Data

The three forest types differ in their climatic preferences. Podocarp forests are predominant on the warmer North Island of New Zealand and appear in more coastal areas at lower elevations and slopes. On average, they are more humid. Beech forests on the other hand are more seasonal (lower isothermality), drier, higher in elevation and cooler. They predominate on New Zealand’s South Island.

Mixed forests lean towards one or the other type. For example, they have a similar elevational profile as podocarp forests, but their mean latitude is closer to that of beech forests. Their temperature preference is naturally between the other types. On average, the three forest types are along a gradient from a subtropical to a temperate forest.

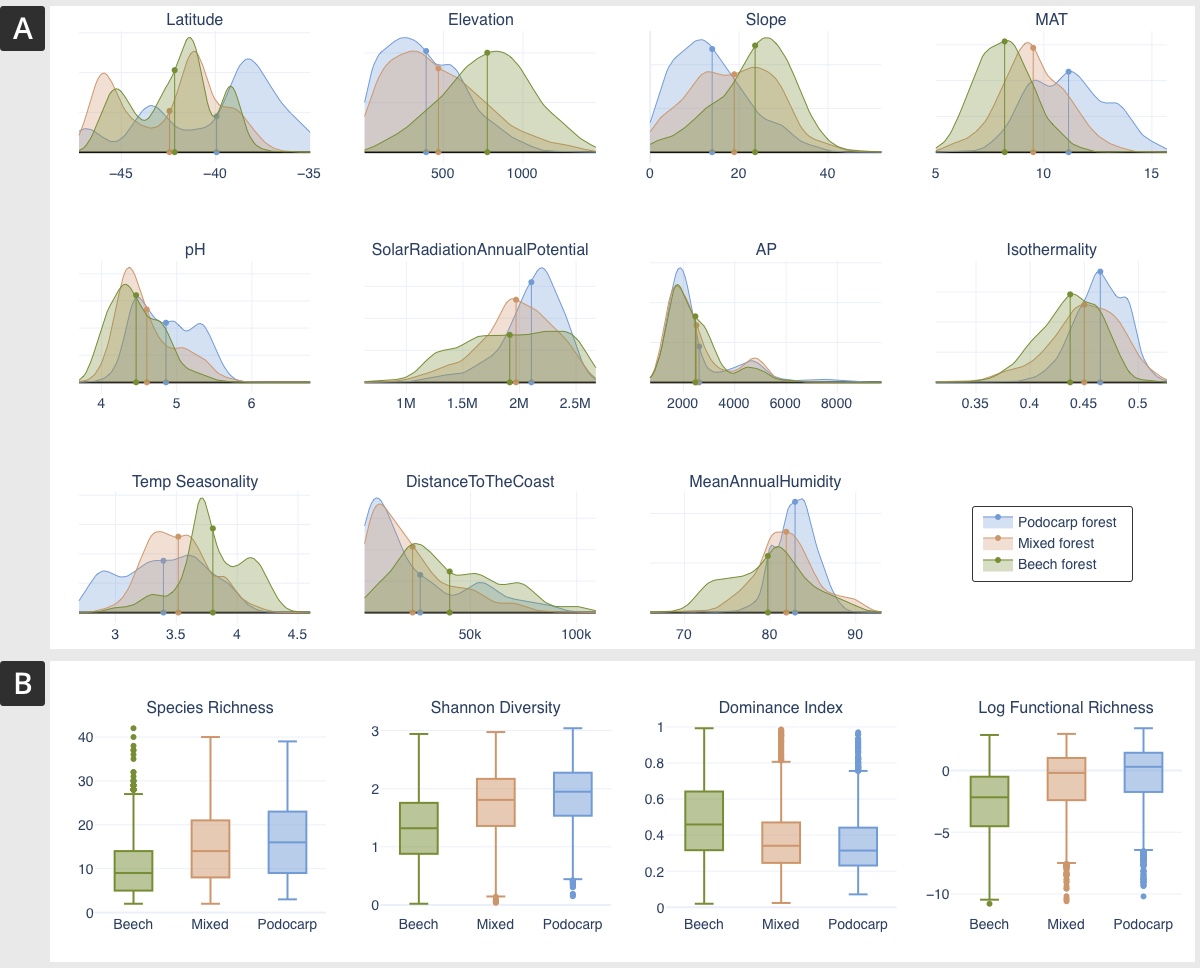

**Figure S1**: Properties of the studied forest types. (A) Distributions of environmental variables across the three forest types. Curves generated via kernel density estimate. Latitude multimodality is due to the three separate islands in New Zealand (North, South, and Stewart Islands). (B) Species richness, Shannon-Weiner Index (measures diversity), Dominance Index (basal cover of dominant species in the plot), and log-transformed functional richness.

### 2 Imputation

In *miceforest* *(Wilson et al., 2023)*, multiple imputation works by iteratively filling in missing values using a random forest algorithm (Breiman, 2001). The basis is a species-trait matrix with each row corresponding to the trait values of a single species. This matrix contains missing values. For each missing value, the algorithm uses the observed data to build a series of decision trees that predict the missing values based on the patterns learned from the complete data. During each iteration, the imputed values are updated by considering both the relationships between variables and the uncertainty associated with the missing data. The algorithm thus exploits correlations between known trait values to impute missing values.

The number of iterations $I$ is a hyperparameter that can positively impact performance, at the expense of runtime. Performance can be further improved by adding phylogenetic or taxonomic information, if available, as a species vector and adding it as additional columns to the species-trait matrix (Debastiani et al., 2021). This is typically done using phylogenetic eigenvectors (Diniz-Filho et al., 1998).

##### 2.1 Taxonomic Eigenvectors

We constructed a taxonomic tree (with equal distances on each branch) for our species using the PhyloT(<https://phylot.biobyte.de/>) website and transformed it into an ordinal distance matrix (i.e., one constructed from a taxonomic tree where all branch lengths are equal to one). This distance matrix was transformed into a set of vectors using phylogenetic eigenvector regression (Diniz-Filho et al., 1998), which yields better imputation results (Penone et al., 2014) and has been successfully utilised previously (Debastiani et al., 2021). The number of available phylogenetic eigenvectors (PVRs) is equal to the number of species. We utilized $E=20$ eigenvectors as this resulted in the best performance.

##### 2.2 Imputation Error

To assess the quality of the imputation and optimize the hyperparameters, we implemented a leave-one-out error scheme:

1. From each trait, we removed a single value (of a random species), leading to $T=7$ additional missing values in the species-trait matrix. This value is denoted by $a_{ti}$ for the trait $t$ and species $i$.
2. We add $E$ PVRs as columns to the amputated species-trait matrix
3. The MICE algorithm imputes missing values, resulting in the previously amputated values $a_{ti}$ now having a value $b_{ti}$. If the imputation were perfect $a_{ti}=b_{ti}$.
4. Steps 1 to 3 are repeated $N$ times until every known trait value is amputated at least once. $N$ in our dataset is therefore the number of species where δ¹⁵N is missing, as this trait has the highest missingness (***Fig. S2*** B).
5. A root mean square error (RMSE) between the imputation value and the true value is computed for each trait $t$ individually and normalised (NRMSE) by the standard deviation of the known trait values across all species $\sigma_{t}$:
   $n{rmse}_{t}=\sqrt{\frac{1}{N}\sum_{i} (b_{ti}-a_{ti})^{2}}\cdot\frac{1}{\sigma_{t}}$

The NRMSE expresses the imputation error in relation to the variation of the trait expressed in %. Since known data is removed to compute the NRMSE, the imputation performance using all available values will be better than when computing the NRMSE. This makes our NRMSE calculation an upper bound of the error.

6. An average across all traits of $n{rmse}_{t}$ yields a single error metric for the entire imputation process.

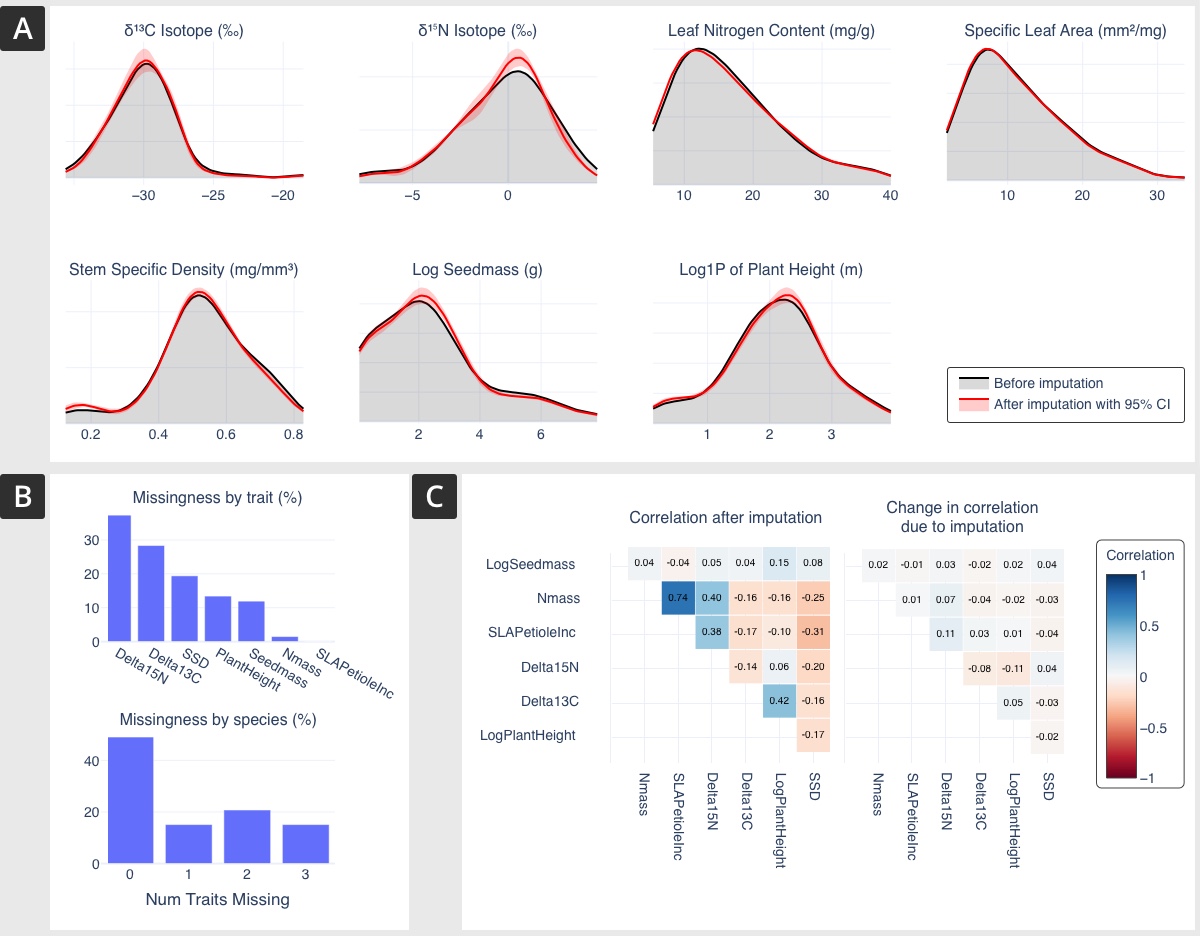

**Figure S2**: Impacts of imputation on trait distributions. (A) Distributions of traits before and after imputation. (B) Missingness. Top: Percentage of species missing a specific trait. Bottom: Percentage of species missing a certain number of traits. (C) Correlation matrix of the traits. Left: After imputation. Right: Difference between the correlation matrix before and after imputation. i.e. changes in correlation induced by the imputation procedure.

##### 2.2 Error and Hyperparameter Choice

The number of MICE iterations $I$ (see (Wilson et al., 2023)) and PVRs $E$ are hyperparameters optimised by exhaustively trying out various combinations (grid search) and comparing the average NRMSE. The lowest mean error across all traits was 10.68% at five iterations and 20 PVRs.

### 3 Supplementary Results

##### 3.1 Changes in climate

Over the last 50 years, mean temperatures have been increasing across all forest types at a rate of ~0.12°C per decade or 0.6°C over 50 years, while seasonality decreased (***Fig. S3***A). Precipitation in all forest types decreased by ~22 mm per decade, corresponding to a decrease of 0.87% of the mean annual precipitation or ~4.8% over the last 50 years. Precipitation seasonality showed a differential response, decreasing in mixed forests and increasing in the other types. Podocarp forests experienced smaller changes in temperature and temperature seasonality, but the strongest increase in precipitation seasonality (***Fig. S3***B).

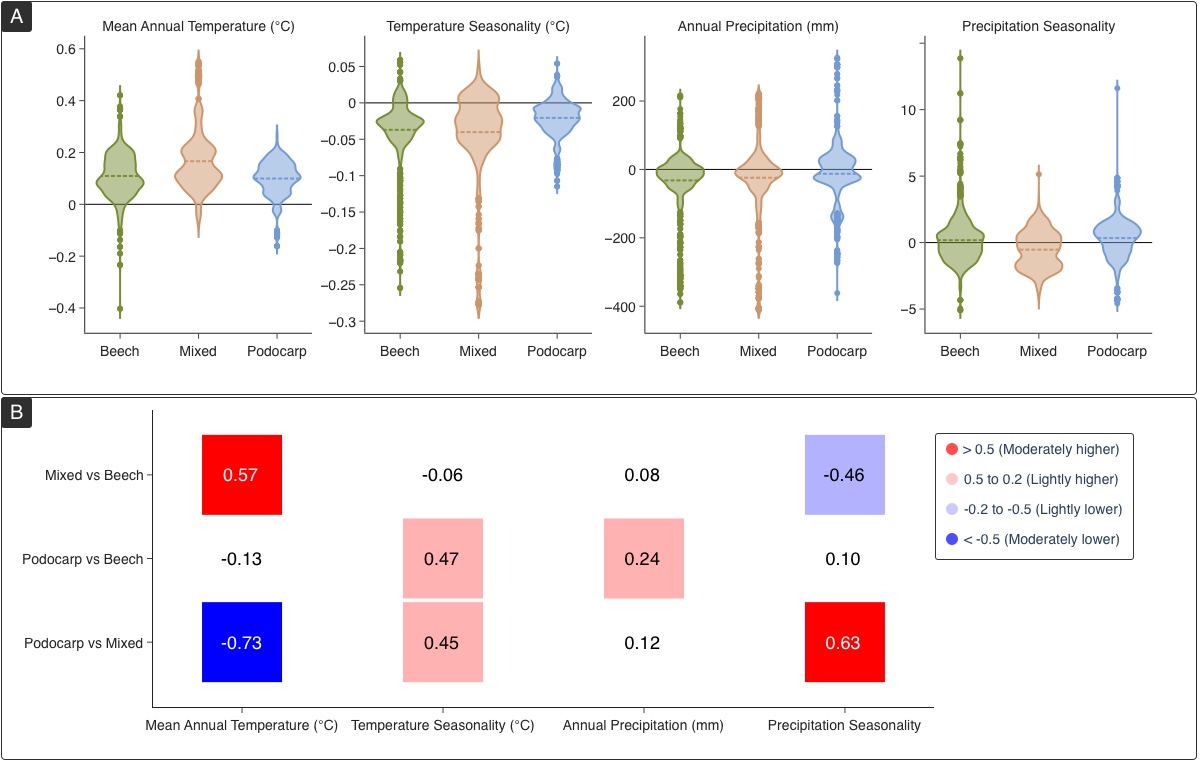

**Figure S3**: Rate of climate change on the three forest types. (A) Changes in temperature, precipitation, and their respective seasonality values per decade. Distribution derived from changes at single forest plots. The dashed line indicates the mean, and the black line is at zero. (B) Differences between distributions in panel (A) were measured by Cohen’s D. Colours represent the strength of difference from moderately lower (darker blue) to moderately higher (darker red).

##### 3.2 Community environmental preferences

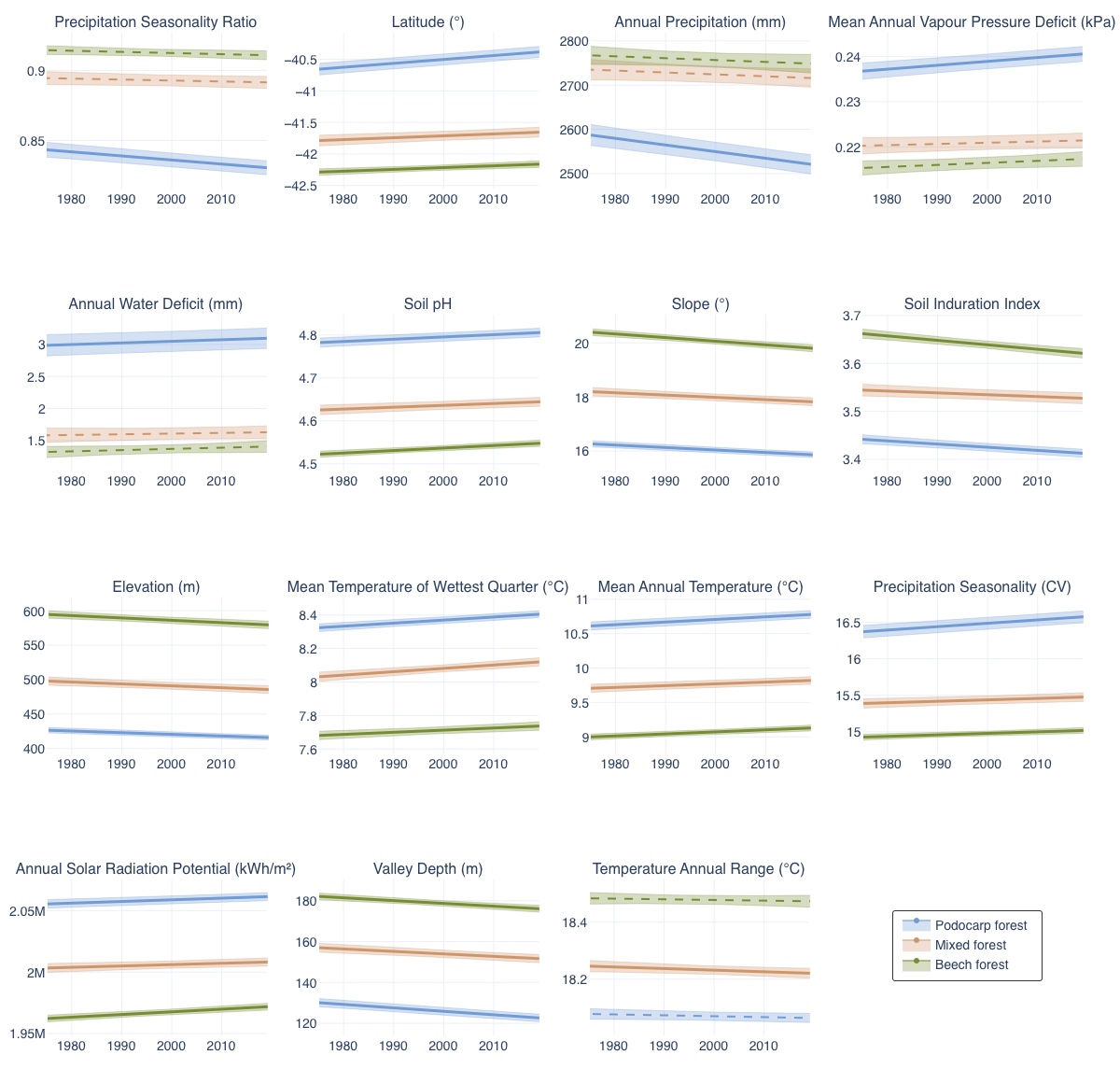

**Figure S4**: Community environmental preferences of all environmental variable traits used in this study over time by forest type. Shaded areas are 95% confidence intervals. Dashed lines imply a non-significant relationship.
